# On the same side: The immune regulatory protein Vista and its ligands interact in cis

**DOI:** 10.1101/2024.08.02.606340

**Authors:** Karina Smorodinsky-Atias, Gil Wiseglass, Mariana Salem, Maya Kashani, Nadir Boni, Alina Artyukhova, Rachel Levy, Rotem Rubinstein

## Abstract

VISTA, an essential immune checkpoint regulatory protein, regulates peripheral T-cell quiescence and tolerance. Despite its potential as a target for anti-tumor and autoimmune disease therapies, uncertainty regarding VISTA’s binding mode and membrane orientation has hindered these developments. Contrary to the prevailing paradigm, we found using cell aggregation assays that VISTA cannot interact with its ligands in *trans* (between cells). Using MST and flow cytometry, we showed that soluble VISTA binds to its ligands, suggesting that VISTA’s membrane orientation restricts *trans* interactions. In contrast, split luciferase complementation assays showed that VISTA interacts with its ligands in *cis*. We propose that a disulfide bond bends VISTA’s Ig domain towards the membrane in an orientation that prevents *trans* while enabling *cis* interactions. Co-expression data analysis from the cancer genome atlas showed a strong correlation between VISTA and its ligand, PSGL-1, consistent with our in-vitro *cis* interaction data. Our findings reveal VISTA’s binding mechanism and suggest an intrinsic inhibition signaling pathway independent of additional cells. Importantly, our experimental framework provides a platform for identifying novel VISTA-targeted therapeutics.

## Introduction

The interactions between immune cell regulatory receptors and their cognate protein ligands play a crucial role in determining the immune response’s strength, duration, and overall quality. Immunotherapies that target these interactions have significantly advanced treatments for cancer, infections, and autoimmune diseases (Wei *et al*, 2018; Baumeister *et al*, 2016; Chattopadhyay *et al*, 2009). The contribution of these protein interaction-focused therapies is evident in anticancer drugs that target Programmed cell death protein 1 (PD-1), PD-Ligand 1 (PD-L1), and Cytotoxic T-lymphocyte associated protein 4 (CTLA-4) among other receptors. However, despite these significant advancements, the limited efficacy of these therapies across different patient populations and cancer types highlights the urgent need to explore alternative immune regulatory pathways (Postow *et al*, 2015; Pardoll, 2012).

A promising candidate for targeted immunotherapy is the regulatory receptor V-domain Ig suppressor of T cell activation (VISTA), a B7 immune regulatory protein family member (Yuan *et al*, 2021; Wang *et al*, 2011; Flies *et al*, 2011). VISTA stands apart from CTLA-4 and PD-1 due to its distinct expression pattern and signaling pathway for immune suppression. As CTLA-4 and PD-1 inhibitory receptors are expressed only on activated T cells, VISTA is constitutively expressed on naïve T cells and has been shown to induce quiescence in these cells (ElTanbouly *et al*, 2020b). Notably, VISTA expression is significantly upregulated following antibody therapy targeting CTLA-4 and PD-1 in prostate and melanoma cancer patients, suggesting its potential role in resistance to these immunotherapies (Gao *et al*, 2017; Kakavand *et al*, 2017). Moreover, the combined blockade of VISTA and PD-1 demonstrates a synergistic anti-tumor response with improved therapeutic efficacy in melanoma and colon cancer mouse models, underscoring the potential benefits of combining immune therapies that target VISTA (Liu *et al*, 2015). However, the development of VISTA-targeted immunotherapies remains impeded by a limited understanding of the interaction mechanisms between VISTA and its ligands.

VISTA is a type I transmembrane protein belonging to the immunoglobulin (Ig) superfamily. The closest, albeit weak, homolog to VISTA is PD-L1, a pivotal immune checkpoint ligand that modulates T cell activity. It has unique sequence and structural features compared to other Ig superfamily members, notably four cysteines not found in other members (Flies *et al*, 2011; Wang *et al*, 2011; Rubinstein *et al*, 2013). These cysteines form two distinctive disulfide bonds, one of them between the A β-strand (the first β-strand in an Ig domain) and the H β-strand (an extra β-strand present in VISTA) (Mehta *et al*, 2019; Slater *et al*, 2020). This distinct feature suggests that VISTA may have unique functional attributes, although the relationship between VISTA’s sequence, structure, and function is not yet understood. One possibility is that this disulfide bond, which bends VISTA’s Ig domain towards the membrane, alters its interaction capability.

VISTA loss of function experiments often result in a dysfunctional immune response, cancer, and autoimmune diseases (Wang *et al*, 2011, 2014; Ceeraz *et al*, 2017; Li *et al*, 2017). Mice with VISTA deficiency or blockade exhibit spontaneous T cell activation, overproduction of inflammatory cytokines, and immune infiltrates in organs such as the lungs, skin, and kidneys. In disease models including encephalomyelitis, hepatitis, asthma, and lupus, VISTA deficiency significantly increases disease incidence and severity (Ceeraz *et al*, 2017; Wang *et al*, 2014; Li *et al*, 2017; Liu *et al*, 2018). Conversely, elevated VISTA expression is associated with poor survival and immunosuppression in human oral squamous cell carcinoma, colon cancer, and gliomas (Wu *et al*, 2017; Deng *et al*, 2019; Ghouzlani *et al*, 2021).

To date, four potential ligands for VISTA have been identified, including V-set and immunoglobulin domain containing 3 (VSIG3) and VSIG8, which have been shown to bind with VISTA and promote T cell inhibition (Chen *et al*, 2022; Wang *et al*, 2019; Yang *et al*, 2017). A third ligand, P-selectin glycoprotein ligand-1 (PSGL-1), has been shown to interact with VISTA in a pH-dependent manner. Under acidic conditions typical of the tumor microenvironment (pH of approximately 6), PSGL-1 binds to a cluster of protonated histidine residues on the surface of VISTA (Johnston *et al*, 2019). Finally, one report proposed that VISTA may possess dual functionality as both a ligand and a receptor through homophylic *trans* interactions (VISTA-VISTA binding between two cells) (Yoon *et al*, 2015). This is particularly intriguing given that VISTA is expressed on both T cells and antigen-presenting cells (APCs) (Yuan *et al*, 2021), suggesting that VISTA could function as its own ligand.

Currently, it remains unclear whether any of the potential VISTA ligands are functional physiologically (ElTanbouly *et al*, 2020a). In addition, VISTA-ligand interaction mechanisms remain unclear, with most models suggesting the interaction occurs between adjacent cells (trans interactions). However, inspired by a recent report on *cis* interaction (on the same cell) between PD-L1 and CD80 on myeloid cells (Chaudhri *et al*, 2018) and noting the functional and expression similarities between VISTA and PSGL-1 on leukocytes, ElTanbouly et al. have proposed that VISTA may interact with PSGL-1 in *cis* rather than in *trans* (ElTanbouly *et al*, 2020a). This uncertainty regarding the precise mechanisms and orientation of VISTA-ligand interactions has hindered progress in developing therapeutics targeting the VISTA signaling pathway.

In this paper, we used cell aggregation assays and found that VISTA and its ligands do not engage via cell-cell *trans* interactions. In contrast, a soluble Ig-fusion version of VISTA could bind to cells expressing PSLG-1 and VSIG3, as shown in flow cytometry analysis. Similarly, using MST, we showed that soluble VISTA interacts with soluble VSIG3, which is consistent with previous studies. These results suggest that VISTA’s orientation on the membrane prevents its interaction in *trans*. Finally, we demonstrated that VISTA could interact with all its potential ligands in *cis* using split luciferase complementation assays. Together, our findings reveal the interaction mechanism for the VISTA signaling pathway.

## Results

### VISTA and its ligands do not engage in trans interactions

Much of the previous evidence for VISTA interactions with its partners was based on soluble forms of at least one of the interacting proteins. To determine whether VISTA can interact in *trans* with its ligands when both are membrane-bound, we utilized cell-aggregation assays. This well-established method is used to study *trans* interactions between receptors and adhesion proteins (Wiseglass & Rubinstein, 2024; Thu *et al*, 2014; Bisogni *et al*, 2018; Schreiner & Weiner, 2010; Wiseglass *et al*, 2024). Our approach involved transfecting two populations of suspension-cultured cells, each expressing either VISTA or one of its ligands, fused to different fluorescent tags. The two cell populations are subsequently mixed and allowed to aggregate for several hours. Binding between VISTA and ligand should induce mixed cell aggregates, which are detectable via fluorescent microscopy.

To minimize artifacts, we conducted two sets of experiments, one using K562 and the other using suspension-adapted HEK293F cells. We mixed cells expressing VISTA C-termini fused to an enhanced green fluorescent protein (EGFP) with cells expressing either VSIG3, VSIG8, or PSGL-1 C-terminally fused to mCherry. For a negative control, we mixed cells expressing EGFP with cells expressing mCherry. For positive control, we mixed cells expressing nectin-1 and nectin-3, which are cell-surface receptors known to interact in *trans* (Satoh-Horikawa *et al*, 2000; Harrison *et al*, 2012). Contrary to our expectation, VISTA and its ligands did not mediate cell aggregation (Figure 1A). This observation, identical to the negative control, indicates *trans* interactions do not occur between VISTA and its ligands. Additionally, cells expressing VISTA did not exhibit homotypic aggregation, indicating that VISTA does not participate in homophilic *trans* interactions. In contrast, the positive control cells expressing Nectin1 and Nectin3 formed large mixed aggregates. These results were consistent across HEK293F and K562 cells (Figure 1A). Because deletion of the cytoplasmic tail was reported to improve protein expression in cell aggregation assays (Schreiner & Weiner, 2010), we tested both wild-type and cytoplasmic-deletion constructs (ΔCyto); however, neither construct led to cell aggregation (Figure 1A and Sup. Figure 1A).

**Figure 1.**
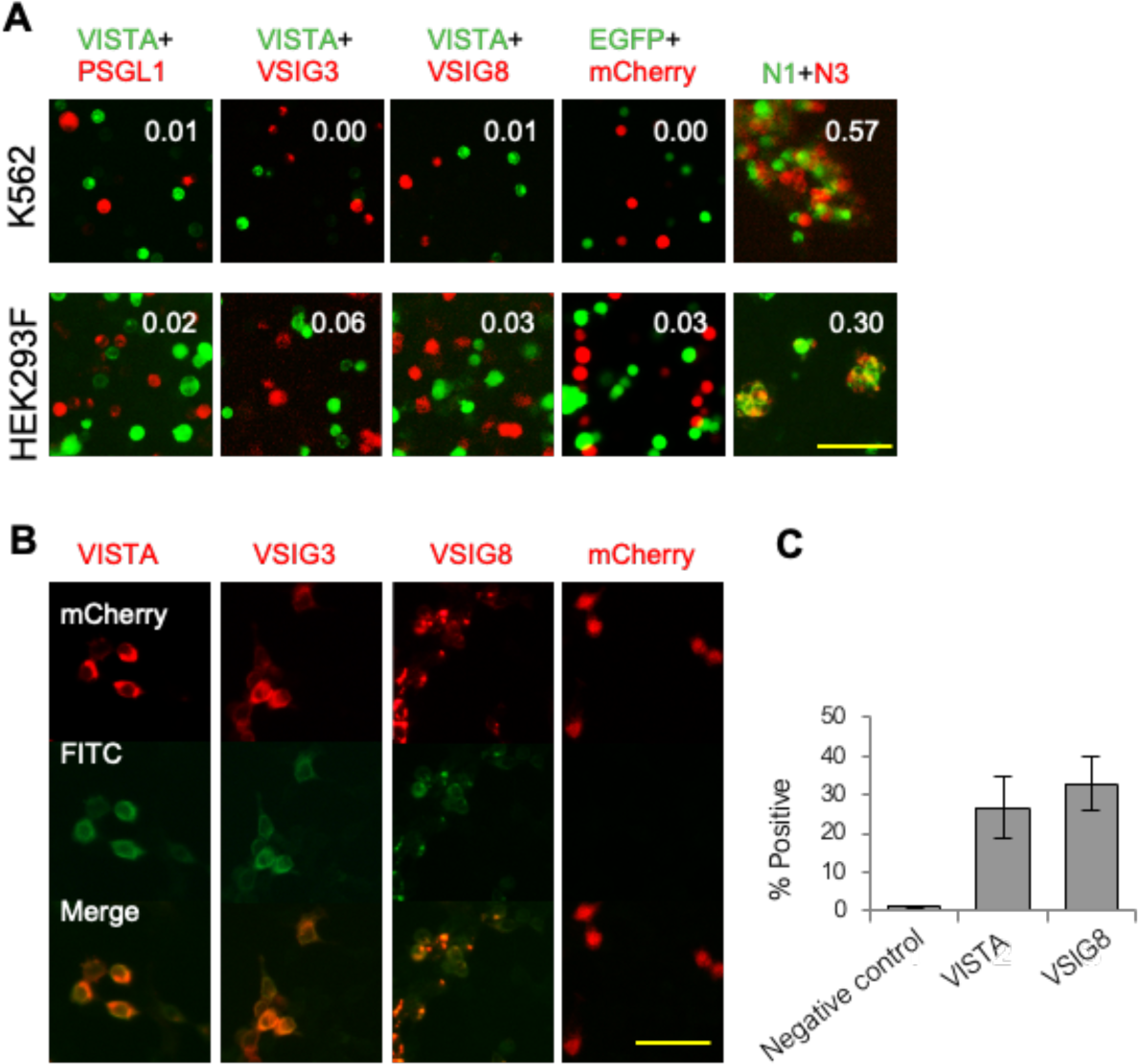
VISTA does not engage in *trans* interactions with its ligands. A) Typical result for cell aggregation assays involving K562 (Left) and HEK293F (Right). Cells expressing differentially tagged VISTA and ligands, or controls (EGFP and mCherry as a negative control, nectin1 and nectin3 as a positive control), are mixed and assayed for aggregation behavior. In both cell types, VISTA and its ligands, as well as the negative control, do not form aggregates, indicating a lack of *trans* interaction. The positive control forms large mixed aggregates, representing strong *trans* binding of the nectins. Scale: 100 µm. B) Immunostaining of the HEK293T cells transiently transfected with VISTA, VSIG3, or VSIG8 cDNAs fused to mCherry fluorescent proteins. Cells were then stained with an anti-HA tag antibody that was detected using FITC-conjugated anti-rabbit antibody and compared to a negative control (mCherry expressing cells). Scale: 50 µm. C) VISTA and VSIG8 are localized on the outer cell surface of K562 cells. K562 cells transfected with VISTA fused to HA tag in the stalk region (VISTA-HA-EGFP) or VSIG8-HA-mCherry were stained with an anti-HA antibody and compared to a negative control (cells expressing wild-type VISTA-EGFP and VSIG8-mCherry) using flow cytometry.

Due to low VSIG3 expression levels in both HEK293F and K562 cells (Figure 1B), we repeated the experiment using NALM6 cells. In these cells, VSIG3-mCherry was strongly expressed and formed homotypic red cell aggregates (Sup. Figure 1B), a result consistent with a previous report of VSIG3 homophilic interaction (Harada *et al*, 2005). In contrast, VISTA did not mediate homotypic aggregates (Sup. Figure 1B) or heterotypic aggregates with VSIG3 expressing cells (Sup. Figure 1B). Overall, these results suggest that VISTA cannot mediate cell-cell interactions with any of its known ligands.

To ensure that the observed lack of *trans* interactions in the cell aggregation assays was not due to inadequate protein cell surface delivery, we performed immunostaining of the transfected cells. HEK293T cells were transfected with either VISTA, VSIG3, or VSIG8, each fused to an HA-tag in the stalk region and stained with HA-tag antibody conjugated to Fluorescein isothiocyanate (FITC). Fluorescent imaging showed a clear staining of the cell membrane, consistent with protein localization on the outer cell surface (Figure 1 B & C).

### Soluble VISTA interacts with ligands

Given the unexpected cell aggregation assay results suggesting VISTA’s inability to bind in *trans*, we sought to confirm its binding capabilities to its ligands in solution, where the cell membrane does not restrict either protein orientation. We selected VSIG3, as its interaction with VISTA had been previously measured in two independent reports (Wang *et al*, 2019; Yang *et al*, 2017). We expressed and purified VISTA and VSIG3 ectodomains and tested their binding by performing a microscale thermophoresis (MST) assay. The MST data clearly demonstrated that VISTA binds to VSIG3 with a disassociation constant (K_d_) of 0.208 ± 0.0795 μM, similar to the previous report (Yang *et al*, 2017) (Figure 2A). To test if soluble VISTA can form homophilic interactions, we conducted an analytical ultracentrifugation (AUC) assay. Sedimentation equilibrium analysis did not reveal measurable homodimer forms of VISTA and was consistent with VISTA behaving as a monomer in solution (Sup. Figure 2B).

**Figure 2.**
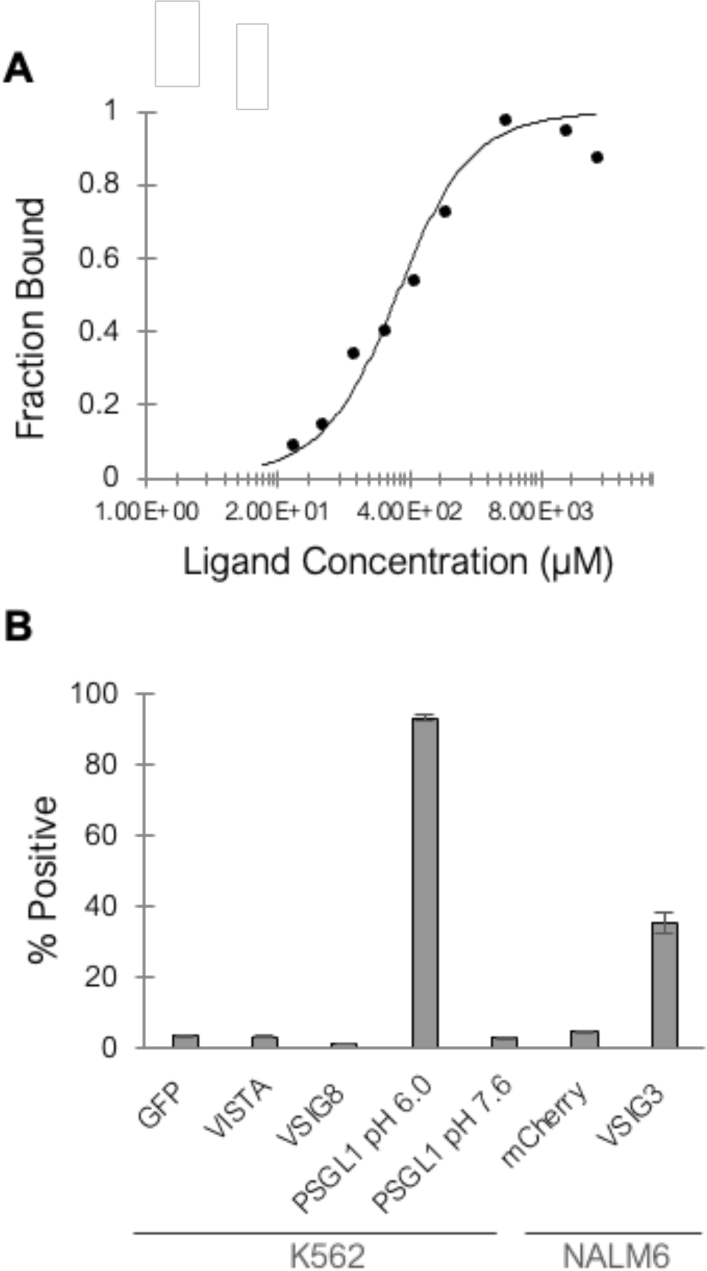
Soluble VISTA binds to soluble and membrane-bound ligands. A) MST binding curve of EGFP fused to the VISTA ectodomain at a concentration of 375 nM in each MST capillary was incubated with variable concentrations of the ectodomain of VSIG3. MST signal was fitted to a single site-binding model yielding the K_d_. B) Flow cytometry analysis shows soluble VISTA-Ig binds to PSGL-1, expressed on K562 cell surface under acidic conditions and with VSIG3 expressed in NALM6 cells. No additional interactions between VISTA and other ligands were detected.

Our investigation into VISTA’s interaction capabilities prompted us to conduct a binding assay between soluble VISTA and its membrane-bound partners. We expressed and purified VISTA-Ig fusion protein and evaluated its binding to cells expressing its ligands by flow cytometry. We did not detect binding of VISTA-Ig to VSIG3, VSIG8, or VISTA expressing K562 cells (Figure 2B). We hypothesized that the lack of binding to VSIG3-expressing cells was due to low expression levels. Indeed, soluble VISTA-Ig did bind to VSIG3-expressing NALM6 cells, which express VSIG3 strongly (Figure 2B). Notably, soluble VISTA exhibited enhanced binding to PSGL-1 in acidic pH compared to binding experiments in physiological pH and no binding to control cells expressing EGFP (Figure 2B). These findings suggest that, while VISTA does not mediate cell-cell interactions in *trans*, it can bind to cells expressing VSIG3 and PSGL-1 when in solution. The lack of binding of VISTA to itself or to VSIG8 was also observed in other studies (Johnston *et al*, 2019; Wang *et al*, 2019). These results underscore the potential influence of membrane constraints on VISTA’s binding capabilities.

### VISTA engages with VSIG3, VSIG8 and PSGL-1 in cis interactions

Many cell-surface receptors function through *cis* interactions with molecules on the same cell membrane (Honig & Shapiro, 2020; Held & Mariuzza, 2011; Hui, 2023). To test whether VISTA interacts with its ligands in *cis*, we used the NanoBiT fragment complementation assays (PCA), adopting a strategy recently reported for studying *cis* interactions between immune regulatory receptors (Chaudhri *et al*, 2018; Garrett-Thomson *et al*, 2020). This assay assesses interactions between two proteins by fusing each protein to a non-functional fragment of a nano-luciferase reporter enzyme. An interaction between the proteins of interest brings together the complementary reporter fragments, enabling detection through the reconstitution of a functional reporter protein (Figure 3A).

**Figure 3.**
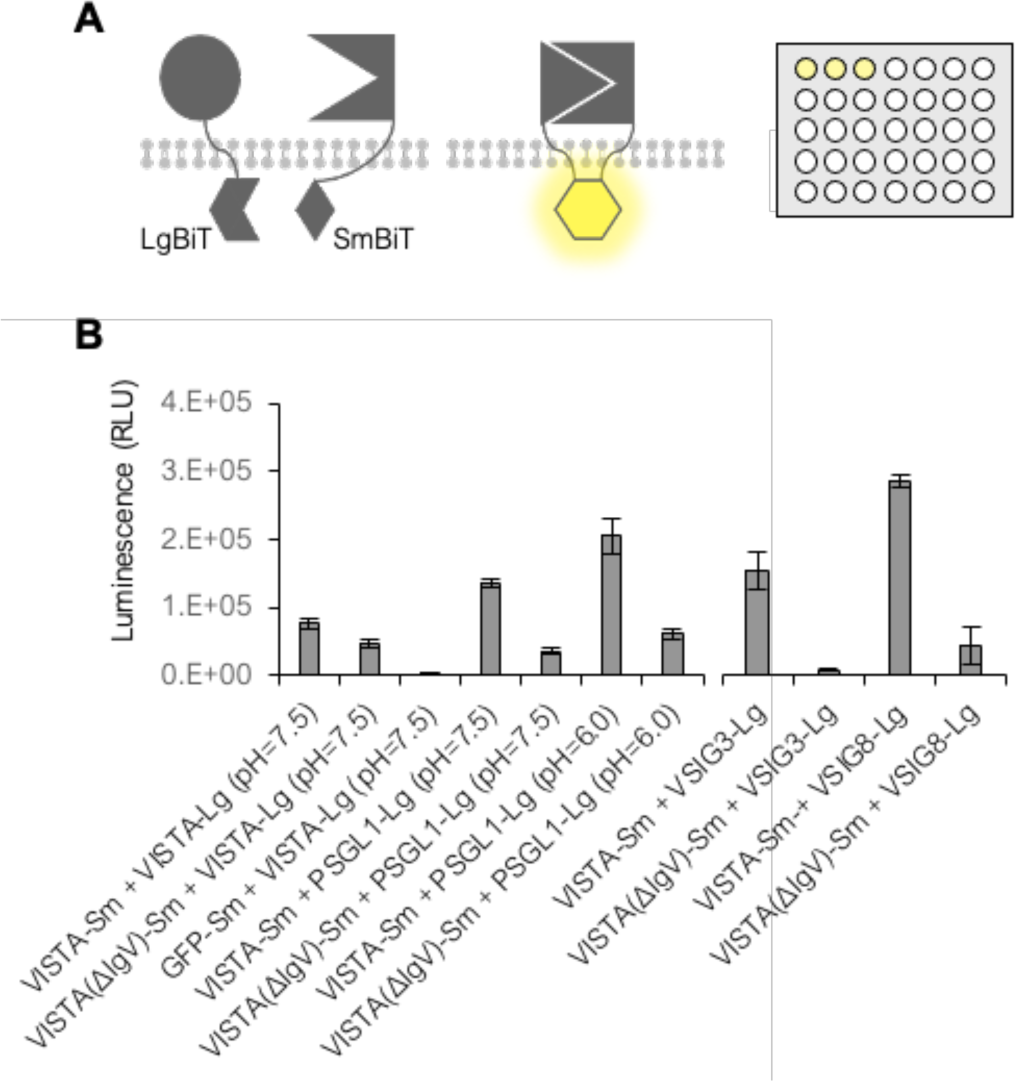
VISTA engages with VSIG3, VSIG8, and PSGL-1 in *cis* interactions. A) An illustration of the NanoBiT assay principle. Proteins of interest are fused to one of two complementary fragments and co-expressed in cells. When both proteins interact in *cis*, the complementary fragments will bind. This binding leads to a detectable luminescence upon application of the NanoGlu substrate. B) HEK293T cells were transiently co-transfected in triplicates in a 96-well plate. NanoGlu substrate was added to each well less than 24 hours after transfection, and luminescence was measured using a BioTek Synergy H1 plate reader. Homophilic *cis* interaction of VISTA as well as heterophilic *cis* interactions between VISTA and VSIG3, VSIG8, or PSGL1 in two different pH were measured. Two distinct experiments are presented in separate panels for clarity.

We designed constructs of VISTA fused to the small fragment of the nano-luciferase enzyme (SmBiT) and constructs of VSIG3, VSIG8, PSGL-1, and VISTA fused to the complementary large fragment of the enzyme (LgBiT). Additionally, we constructed a negative control by fusing EGFP to the LgBiT terminus. Co-transfection of VISTA-SmBiT with any one of the VSIG3-, VSIG8-, and PSGL-1-LgBiT constructs led to nano-luciferase reconstitution, apparent by the strong luminescent signal in the presence of NanoGlu substrate (Figure 3B). In contrast, co-transfection with VISTA-SmBiT and the negative control EGFP-LgBiT result in minimal luminescence (Figure 3B). These results indicate that VISTA can engage in *cis*-homophilic as well as heterophilic interactions with VSIG3 and VSIG8.

Next, we aimed to determine whether the observed *cis* interactions involved VISTA’s Ig domain, which mediates its binding to protein partners (Johnston *et al*, 2019; Xie *et al*, 2021). To this end, we constructed VISTA lacking its Ig domain (VISTAΔIg) fused to the SmBiT fragment (VISTAΔIg-SmBiT). We observed that co-expression of VISTAΔIg-SmBiT with VSIG3-LgBiT, VSIG8-LgBiT, or PSGL-1-LgBit resulted in a significant decrease, by several orders of magnitude, in the luminescent signal compared to the wild-type VISTA, supporting the essential role of the Ig domain in VISTA’s *cis* binding to VSIG3, VSIG8, and PSGL-1 (Figure 3B). Intriguingly, co-expression of VISTAΔIg-SmBiT with VISTA-LgBiT resulted in a diminished luminescent signal compared to co-expression of wild-type VISTA, though not as significant as with heterophilic interactions. Thus, we could not detect Ig-dependent homophilic *cis* interaction of VISTA (Figure 3B). It is possible that VISTA can form homophilic *cis* interactions via the transmembrane or cytoplasmic domains of the protein. The co-transfection of HEK293T cells with VISTA-SmBiT and PSGL-1-LgBiT demonstrated high luminescence at pH 6.0. Remarkably, a similar experiment at pH 7.5 resulted in significantly reduced luminescence, demonstrating that our assay captured VISTA-PSGL-1 pH-dependent *cis* interaction (Figure 3B).

*cis*-interacting proteins are expected to have similar expression patterns, a characteristic that was previously leveraged to identify functional and regulatory networks (Stuart *et al*, 2003; Wang *et al*, 2016). We applied a similar strategy using the TCGA Pan-Cancer Atlas database (Cancer Genome Atlas Research Network *et al*, 2013) which contains mRNA expression data from various cancer tissue samples (see Methods). Figure 4 shows the co-expression patterns of VISTA with either VSIG8, VSIG3, PSGL-1, or PD-L1 as a positive control. A small correlation between VISTA and PD-L1 mRNA expression was observed, consistent with their known overexpression in multiple cancer types (He *et al*, 2021; Rezouki *et al*, 2023). A strong correlation was found between VISTA and PSGL-1 across various cancer types (Figure 4D and SI Figure 3), suggesting a functional connection and supporting a *cis* interaction-dependent relationship between VISTA and PSGL-1. In contrast, no correlation was observed between VISTA and VSIG8 or VSIG3 (Figure 4 A-C), which may suggest either the absence of a linear functional relationship or non-significant physiological functionality in cancer cells.

**Figure 4.**
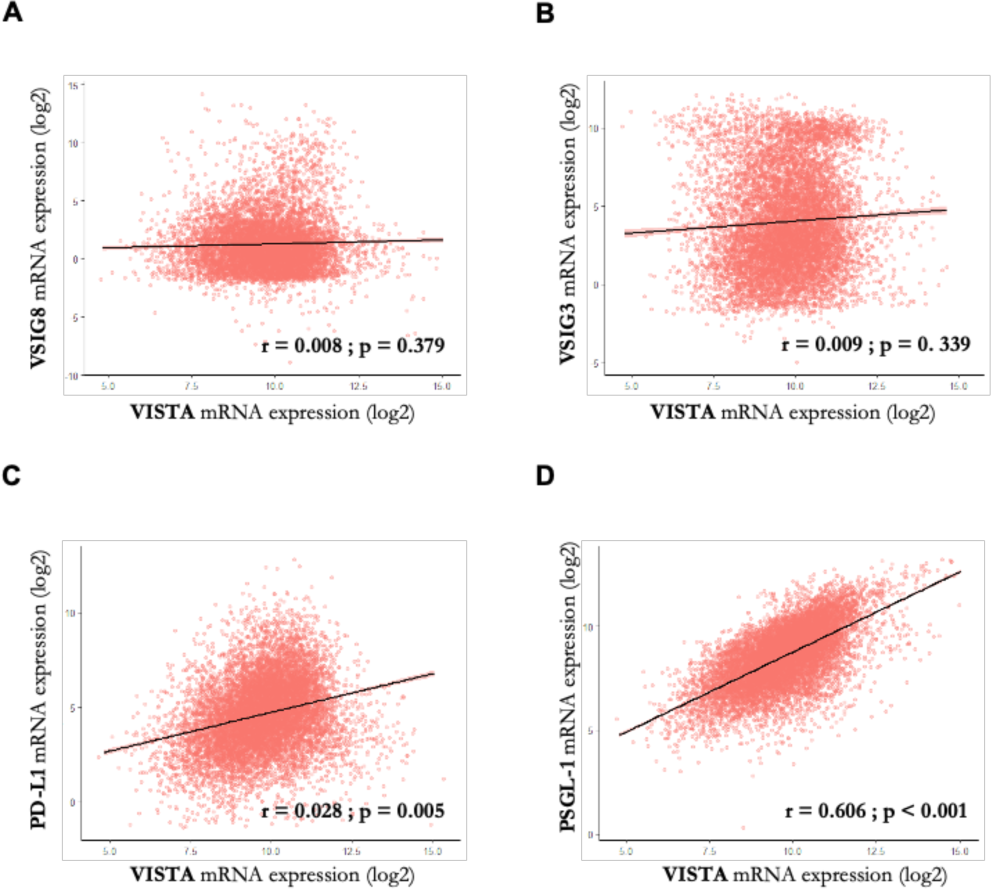
VISTA expression pattern correlates with PSGL-1 in various cancer types. Co-expression of VISTA with VSIG8 (A), VSIG3 (B), PD-L1 (C), and PSGL-1 (D). The plots depict the normalized mRNA expression across various cancer types and samples (data from TCGA, see Methods section). A clear correlation is only observed between VISTA and PSGL-1 (D).

## Discussion

Using cell aggregation assays, flow cytometry, MST, AUC, and split luciferase complementation assays, we show that VISTA binds to its ligands in *cis* but not in *trans*. Our data support the notion that this behavior is due to spatial constraints and orientation of VISTA on the cell membrane, which hinder VISTA’s ability to interact with ligands on neighboring cells.

While the current view of VISTA activation involves *trans* interactions, this is likely because most immune regulatory interactions have been usually studied in the context of *trans* interactions. A close examination of the VISTA literature reveals no direct evidence for *trans* interaction between VISTA with its currently known ligands. The various studies of VISTA interactions have explored the interactions between VISTA and its ligands and their impact on immune function only when either VISTA or its ligands were in soluble form (Chen *et al*, 2022; Wang *et al*, 2019; Yang *et al*, 2017)(Johnston *et al*, 2019). Here we utilized cell aggregation assay as a method to detect interactions between proteins specifically when both parties are membrane-bound. Our results have consistently demonstrated a lack of *trans* interaction between VISTA and any of its currently known ligands.

To determine if membrane constraints were preventing interactions, we tested whether soluble VISTA could bind to its ligands. We performed an MST assay using purified VISTA and VSIG3 ectodomains, confirming binding in solution with a K_d_ in the micromolar range, typical for heterophilic and homophylic interactions of other Ig superfamily immune regulatory proteins such as PD-1:PD-L1 and CRTAM, among others (Zhang *et al*, 2004; Rubinstein *et al*, 2013; Davis *et al*, 2003). Additionally, findings from flow cytometry binding assays showed that soluble VISTA interacts with a membrane-bound PSGL-1, particularly under acidic conditions that mimic the tumor microenvironment. These findings indicate that VISTA can, in fact, interact with its ligands when it is not constrained by the cell membrane, which supports our hypothesis that the membrane-bound orientation affects VISTA’s *trans* binding abilities. Notably, evidence that VISTA can be cleaved from the cell membrane and consequently interact with cells (Yasinska *et al*, 2020) suggest that an interaction between soluble VISTA and membrane bound ligands could be physiological relevant.

NanoBiT fragment complementation assays demonstrated that VISTA engages in *cis* interactions with its ligands VSIG3, VSIG8, and PSGL-1. These *cis* interactions are mediated via VISTA’s Ig domain, as evidenced by the significantly reduced luminescent signal observed when this domain is deleted. Interestingly, despite previous reports of VISTA self-interaction (i.e., VISTA with VISTA as its ligand) (Yoon *et al*, 2015), we did not detect this behavior in our assays. Thus, in solution, an AUC analysis revealed that VISTA ectodomains behave as a monomer. Moreover, VISTA did not engage in homophilic *trans* interaction in our cell aggregation assay. Lastly, we did not detect Ig-dependent *cis*-homophilic interaction, although it is still possible that VISTA can form *cis* interactions via its transmembrane and cytoplasmic domains.

Our results suggest that VISTA interacts in *cis* but does not interact in *trans* due to geometrical constraints involving the protein and the membrane. One likely mechanism creating this constraint is a disulfide bond, unique to VISTA, that likely bends its Ig domain toward the membrane and hinders *trans* interaction with its ligands. This structure differs from that of the typical immune regulatory protein, where the N-terminal Ig domain extends straight from the membrane, exposing its interface for *trans* interactions (Chattopadhyay *et al*, 2009). The functional importance of the VISTA-specific disulfide bond was demonstrated by its deletion, which resulted in the loss of VISTA’s inhibitory function (Slater *et al*, 2020). Interestingly, structural modeling of VISTA’s interactions with its ligands - based on its closest homolog PD-L1 binding with PD-1 - suggests that this domain bending does not prevent *cis* interactions with its ligands (Figure 5A-B).

**Figure 5.**
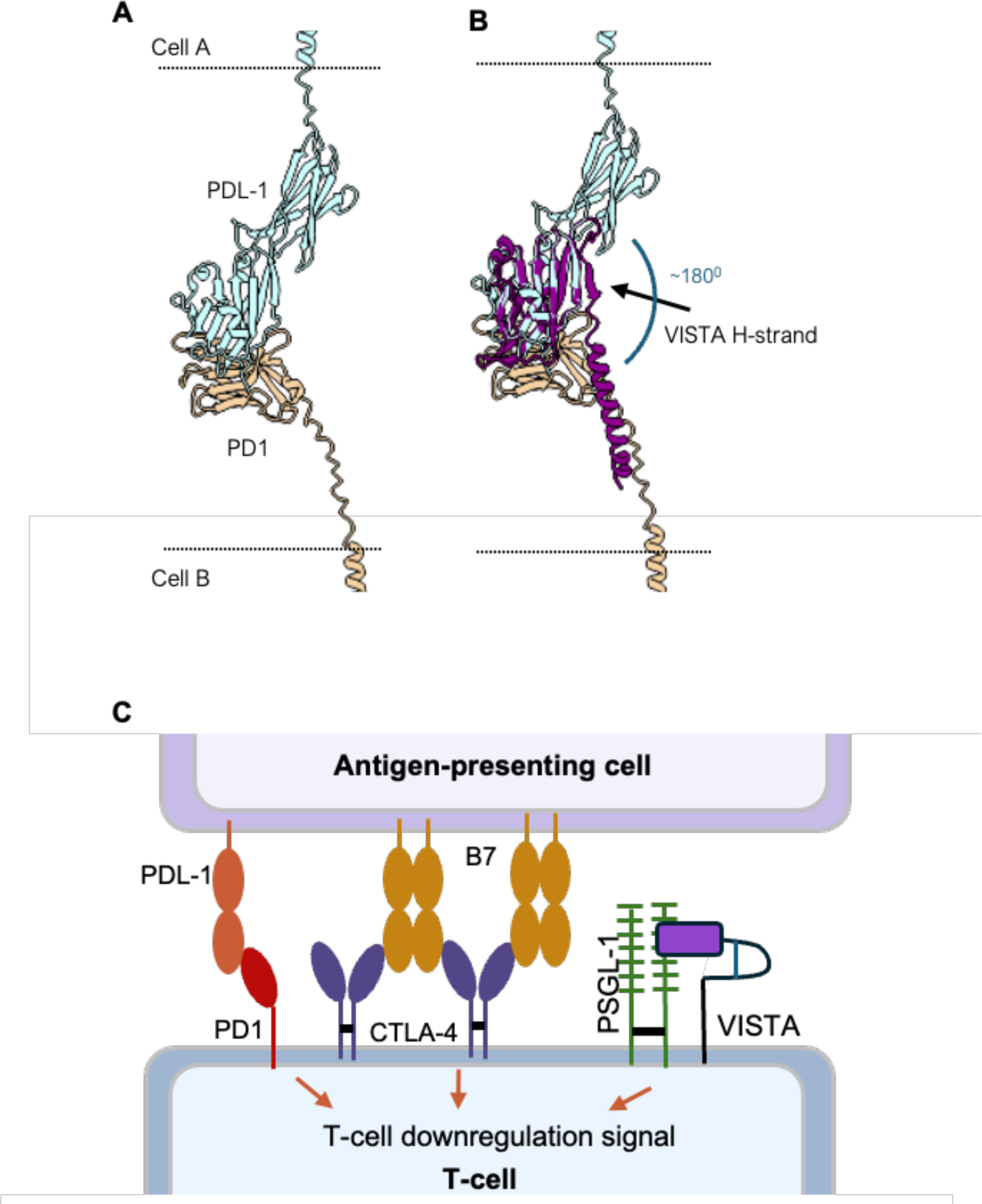
A-B) A comparison of VISTA’s orientation with PD1:PDL-1 complex. A) Structural model of the complex formed between PD1 (tan color) with PDL-1 (light blue, PDB code 3BIK). The stocks and single-pass membrane helices are modeled only for presentation purposes (not from the PDB). Cell membranes are depicted as black dots. B) Superposition of the VISTA Ig domain (in purple, PDB code 6OIL) on PDL-1 N-terminal Ig domain. The VISTA H strand is bent causing an approximately 180 degrees shift in the position of its C-terminal transmembrane domain. Assuming a homologous interface with PDL-1, this orientation will favor a *cis* rather than a *trans* interaction. C) Schematic representation of immune inhibitory checkpoint proteins interactions. CTLA-4 and PD-1 on T-cells engage in *trans* interactions with B7 and PD-L1 on APC cells, triggering T-cell downregulation signaling pathways. In contrast, our in-vitro data suggests that *cis*-complexes between VISTA and its ligands, including PSGL-1, promote T-cell intrinsic inhibitory signal without interacting with other cells.

While *trans* interactions between immune regulatory receptors are critical in modulating the immune response, recent studies have revealed the importance of *cis* interactions in immune regulation (Hui, 2023). The combination of *cis* and *trans* interactions between receptors and ligands, could compete and thereby refine the *trans* binding specificity and requirement for receptor activation. In cases where *cis* and *trans* interactions are mediated by an overlapping interface, *cis* interaction can mask the *trans* interface and prevent spurious non-specific interactions. Such mechanisms have been suggested for sidekick, HVEM:BTLA, PDL1:CD80, and semaphorin:plexin, for example (Goodman *et al*, 2016; Chaudhri *et al*, 2018; Garrett-Thomson *et al*, 2020; Oh *et al*, 2020; Sugiura *et al*, 2019; Rozbesky *et al*, 2020). *cis* interactions between receptors can also generate a complex combinatorial cell-cell recognition that depend on the *trans* specificity of every protein involved, as in the case of clustered protocadherins (Wiseglass *et al*, 2024). Importantly, there are compelling evidence that *cis* interactions can directly activate receptors, for example, Notch interactions (Sprinzak *et al*, 2010), CD2 and its ligands (Li *et al*, 2022), and CD28 (Zhao *et al*, 2023).

Our results suggest that VISTA interacts in *cis* with its ligands. Assuming the function of VISTA as a mediator of quiescence of T-cells (ElTanbouly *et al*, 2020b), one intriguing implication for VISTA *cis* interaction on its immune function could be that these *cis* interactions generate an intrinsic T-cell inhibition signal. This signal would maintain naïve T-cells in a quiescence state, independent of an external cell-cell interaction (Figure 5C). In this context, the NanoBiT fragment complementation assays employed here could be used to identify molecular inhibitors and enhancers of VISTA *cis* interactions. These inhibitors could serve as potential drugs for cancer treatment by enhancing the immune response. Conversely, molecular enhancers that strengthen VISTA’s *cis* interactions and likely promote immune inhibition might be developed as potential therapies for autoimmune diseases.

## Material and Methods

### Plasmids and cDNAs

Human VISTA, VSIG3, and VSIG8 cDNAs were cloned into pcDNA3.1 plasmids with a C-terminal EGFP or mCherry fluorescent tag. These plasmids were used for performing cell-aggregation assays, as well as initial templates for cloning the various truncated and mutated constructs. For protein expression, the same cDNAs were cloned into a paSHP-H mammalian expression plasmid containing a N’-terminal Bip signal sequence and a C-terminal 8x Histidine tag to facilitate protein purification (π-Bip). For the NanoBiT assays, sequences coding for the Large BiT (LgBiT, 159 residues) and Small BiT (SmBiT, 11 residues) fragments were introduced C-terminally to the VISTA, VSIG3 and VSIG8 cDNAs in the pcDNA3 plasmids after a 14-18 residue long Gly-Ser-Arg linker.

### Cell lines and culture media

HEK293T cells were cultured in DMEM (Sartorius #01-055-1A) supplement with 10% FBS (Sigma-Aldrich® #F7524), GlutaMAX™ Supplement (Thermo-Fisher #35050038), sodium pyruvate (Sartorius #03-042-1B) and Pen/Strep solution (Sartorius #03-031-5B), at 37°C and 5% CO_2_. For subculturing, cells were washed with PBSx1 (Sartorius #02-023-5A) and detached from the vessel by incubation with Trypsin C solution (Sartorius #03-053-1A). FreeStyle™ 293F Cells (Thermo-Fisher #R79007) were cultured in FreeStyle™ 293 Expression Medium (Thermo-Fisher #12338026) and grown at 37°C with 8% CO_2_, at constant 135 rpm orbital shaking. K562 cells (ATCC #CCL-243™) were cultured in IMDM (Sartorius #01-058-1A) supplement with 10% FBS (Sigma-Aldrich® #F7524) and Pen/Strep solution (Sartorius #03-031-5B), at 37°C and 5% CO_2_. NALM6 cells (ATCC #CRL-3273™) were cultured in RPMI (Sartorius #01-103-1A) supplement with 10% FBS (Sigma-Aldrich® #F7524) and Pen/Strep solution (Sartorius #03-031-5B), at 37°C and 5% CO2.

### Transfection and protein purification

HEK293T transfections were performed using calcium-phosphate as described in (Kingston *et al*, 2001). HEK293T cells were seeded one day preceding transfection, aiming for 70% cell confluency. Two hours prior to the transfection, the media was replaced to make sure that the pH is in the optimal range. Transfection mixture was prepared by gradually adding DNA dissolved in 250 mM calcium chloride to an equal volume of HEPES buffered solution and incubated for 20 minutes at RT allowing the formation of a fine precipitate. The mixture was added dropwise to the culture. Day after transfection, cells were washed twice with PBSx1 and fed with complete DMEM.

FreeStyle™ 293F cells grown in serum-free media (Invitrogen) were transfected using polyethyleneimine PEI MAX® reagent (Polysciences #24765) according to manufacturer’s instructions. The following morning, fresh media was supplemented, and the cells were cultured for up to 6 days until protein harvest. Secreted proteins were purified using nickel-nitrilotriacetic acid (Ni-NTA) affinity chromatography. Proteins for the MST and AUC analysis were further purified using size exclusion chromatography over Superdex 200 26/60 column (Cytiva) on an AKTA pure fast protein liquid chromatography system (Cytiva).

For lentivirus production, HEK293T cells were seeded in T-75 dishes (Greiner #658170) and co-transfected by calcium phosphate method (Kingston *et al*, 2001) with 3rd generation lentiviral packaging plasmids VSV-G, pLP1 and pLP2 along with a pLV-CMV lentiviral vector carrying a EGFP- or mCherry-fused protein. The lentiviral particles were harvested along with the media 24 and 48 hours after transfection. The media was filtered using a 0.45 µm filter (Millipore #SLHV033R) and concentrated.

For lentiviral infection, 1×10^6^ NALM6 cells were mixed with the concentrated media supplemented with 8 µg/ml Polybrene (Sigma-Aldrich® #TR-1003-G) in a 15-ml conical tube, and spinoculated for 60 minutes at 32°C and 800 xg. Following infection, the sup was removed, and the cells resuspended in 2.5 ml fresh RPMI media and placed in a CO2 incubator. After five-day incubation, the infection process was repeated for a total of two infection cycles.

### Cell aggregation assay

Cell aggregation assays were performed using K562 and HEK293F cells as described in (Wiseglass *et al*, 2024; Wiseglass & Rubinstein, 2024). Mixing quantification was performed using the CoAg index FIJI plugin (Bisogni *et al*, 2024). Specifically, for the K562 assays: 1×10^6^ cells were transfected with 10 µg plasmid DNA using electroporation. The following morning, 250 µl of cells from each population were mixed in 6-well plates with 1.5 ml of complete DMEM. Plates were incubated (37°C, 5% CO2) on a rocking platform at 15 RPM for 3 hours, then captured using Nikon Eclipse Ts2 inverted microscope. For the HEK293F assays: 2 ml of FreeStyle™ 293-F cells in density of 1×10^6^ cells/ml were transfected with 1.25 µg plasmid DNA using PEI MAX® (49553-93-7). Three hours post transfection 1 ml of cells from each population were mixed in 6-well plates. Images were captured two days post-transfection. And for the NALM6 assays: NALM6 cells were infected with the indicated constructs fused to GFP or mCherry fluorescent proteins. 2.5×105 infected NALM6 cells from each transfection population were mixed in an un-treated 6-well plate (SPL life-sciences #32006) and were allowed to aggregate for up to 24 hours inside a CO2 incubator. Imaging of the aggregates was performed using Nikon ECLIPSE Ts2 inverted microscope.

### Surface expression quantification

K562 cells were transfected with either VISTA-HAΔCyto or VSIG8-HAΔCyto plasmids (HA sequence: YPYDVPDYA). The cells were immunostained with Anti-HA Rabbit Monoclonal Antibody (Cell Signaling #3724S) at a dilution of 1:1000 in PBSx1 (Sartorius #02-023-5A) with 1% FBS (Sigma-Aldrich® #F7524) for 45 minutes. Following the primary staining, the cells were incubated with goat Anti-Rabbit IgG H&L (Alexa Fluor® 647) secondary antibody (ab150079) at a dilution of 1:1000 for 45 minutes. Fluorescence was quantified using the S1000EXi flow cytometer, allowing for the assessment of the tagged proteins expression levels.

### Flow cytometry binding experiments

HEK293F cells (1×10^6^ cells/ml, a total of 50×10^6^) were transfected with 31.25 µg π-Bip-VISTA(ΔCyto)-Fc-His plasmid (transfection and purification protocol are described in the “Recombinant protein expression and purification” methods section. To prepare the purified protein for subsequent assay, the buffer was replaced to PBSx1 (Sartorius #02-023-5A) or PBSx1 adjusted to pH 6.0 using HCl. The protein was concentrated to approximately 0.3 mg/ml and then supplemented with 1% FBS (Sigma-Aldrich® #F7524). K562 cells transfected with either pcDNA3-VISTAΔCyto-EGFP, pcDNA3-VSIG3ΔCyto-mCherry, pcDNA3-VSIG8ΔCyto-mCherry, pcDNA3-PSGL-1ΔCyto-mCherry or pcDNA3-EGFP (negative control) were prepared for immunostaining as described in the “Surface expression quantification” methods section, with the exception that soluble VISTA was used as the primary antibody and incubated with the cells for 30 minutes at 37°C.

### Surface expression immunostaining

HEK293T cells were transfected with either VISTA-HA(FL), VSIG3-HAΔCyto, VSIG8-HAΔCyto or mCherry (negative control) plasmids. The cells were fixed with 4% paraformaldehyde and permeabilized with methanol. After washing and blocking the cells with 1% BSA in PBSX1 the cells were incubated with Anti-HA Rabbit Monoclonal Antibody (Cell Signaling #3724S) at a dilution of 1:1000 in PBSx1 (Sartorius #02-023-5A) #F7524) for 1 hour. Following the primary staining, the cells were incubated with Goat Anti-Rabbit IgG Antibody, FITC conjugate secondary antibody (Sigma-Aldrich® #AP187F) at a dilution of 1:1000 for 1 hour. Fluorescence imaging was performed using Nikon ECLIPSE Ts2 inverted microscope.

### NanoBiT

HEK293T cells were seeded in a white 96-well plate (SPL life-sciences #30197) one day before transfection. 0.2 μg from each of the two NanoBiT constructs were used for the calcium-phosphate mediated transfection (see above). 24 hours post transfection, the media was replaced to 0.1 ml Opti-MEM^TM^ and a fresh Nano-Glo® Live cell Reagent (Promega #N2011) was prepared in a 1:20 dilution. 25 μl of reagent were added to each well, and the plate was gently mixed for 5 minutes. Immediately after mixing luminescence was measured every 10 minutes for a 70 minutes period using a BioTek Synergy H1 plate reader.

### Analytical ultracentrifugation sedimentation velocity experiments

Sedimentation velocity experiments were performed at 20°C utilizing a Beckman-Coulter XL-I analytical ultracentrifuge instrument, six-sector cells, and an An60Ti rotor. Protein buffer composition was 150 mM NaCl and 40 mM HEPES buffered saline (HBS) pH=8.0. Absorption scans were collected at 280 nm, rotor speed of 50,000 rpm, and at three protein concentrations (5, 15, and 30 µM). Protein concentration was estimated from the extinction coefficient determined using the ProtParam webserver (Gasteiger *et al*, 2005). Buffer density viscosity and protein partial specific volume were calculated using SEDNTERP software (Philo, 2023).Data was analyzed using the SEDFIT software (Schuck, 2000) to calculate the sedimentation coefficient distributions.

### mRNA co-expression analysis

mRNA expression levels of VISTA and its ligands were analyzed using the TCGA PanCancer Atlas database via cBioPortal (http://www.cbioportal.org/) (Cerami *et al*, 2012; Gao *et al*, 2013; de Bruijn *et al*, 2023). This database includes normalized RNA-seq from 10,071 samples across 32 cancer types. The mRNA co-expression between VISTA to VSIG3, VSIG8, PD-L1, and PSGL-1 was evaluated by Pearson’s correlation coefficient using R version 4.1.2 (R, 2021).

### MST

To measure the binding of VISTA ectodomain to VSIG3 ectodomain, VISTA was expressed fused to EGFP. We then measured MST binding curves by incubating the fluorescently labeled VISTA with VSIG3 in dose dependent manner using the Monolith NT.115 instrument. Proteins buffer composition was 150mM NaCl with 40mM HEPES pH=8.0. Two series of 16 MST measurements were performed in 16 capillaries.

## Acknowledgments

We are grateful to Dr. Iris Ben-Dror and Prof. Joel Hirsch for their help with the analytical ultracentrifugation, Dr. Amit Kessel for insightful discussions and Prof. David Sprinzak for valuable comments on the manuscript. This work was supported by grants from the Israel Cancer Research Fund (ICRF 19-203-RCDA to R.R.) and the Israel Science Foundation (1463/19 to R.R).

## Author contributions

R.R. conceptualized the study, K.S.A., G.W., M.S., M.K., A.A., and R.L. performed data collection, R.R., K.S.A., G.W., M.S., and N.B. analysis and interpretation of results, and R.R., K.S.A, G.W, M.S. and N.B manuscript preparation.

